# Inherent Dynamics of Maltose Binding Protein (MBP) are Immune to the Native Environment

**DOI:** 10.1101/2022.03.15.484495

**Authors:** Johannes Thoma, Björn M. Burmann

## Abstract

Biophysical characterizations of proteins typically rely on a reductionistic approach, studying proteins in a highly purified from and in absence of their natural cellular environment. Little is known about how the highly crowded conditions prevalent within living cells influence the dynamic structures proteins on the molecular level. To address this outstanding question, we characterize here the dynamic behavior of the periplasmic model protein MBP from *Escherichia coli in situ*, confined in the native lumen of bacterial outer membrane vesicles. To this end we determine the dynamics of side-chain methyl groups of MBP across several timescales and compare them to purified *in vitro* MBP. We find that the inherent dynamics of MBP are surprisingly insensitive to the native cellular environment and that the molecular motion of the protein is mainly impacted on a global level.

---

It is a long-standing question how cellular conditions influence proteins on the molecular level, structurally as well as dynamically (1, 2). This question is particularly prevalent against a backdrop of protein studies, which traditionally characterize proteins purified to homogeneity and “free of all other biomolecules” (3). While decades of biophysics without doubt produced amazing insight into the inner molecular workings of proteins, their behavior within cellular environments is only beginning to be understood, not least due to the lack of suitable methods.

Within cells proteins are in constant direct interaction with a plethora of other proteins, solutes, and biological surfaces under highly crowded conditions. One particularly crowded cellular environment is the periplasmic space of Gram-negative bacteria, which contains proteins and other solutes at extraordinarily high concentration, exceeding 300 mg/ml (4). We recently established a method to utilize outer membrane vesicles (OMVs) from *Escherichia coli*, enriched luminally with selected proteins, for the structural characterization of periplasmic proteins in their native environment using high-resolution solution NMR spectroscopy (5). Compared to existing in-cell NMR methods these vesicles come with strongly reduced background signals, due to the missing bulk of cytosolic proteins, and their long-term stability allows recording detailed multidimensional NMR experiments on long timeframes, making OMVs particularly suited to characterize periplasmic proteins.

Here, we enriched the well-characterized model protein MBP in the lumen of OMVs to compare the dynamical behavior of MBP between the native cellular environment and the purified state. To this end we utilized the highly sensitive methyl-TROSY NMR methodology (6), based on selectively labeled side-chain methyl groups of isoleucine, leucine and valine (ILV, see Fig. S1) residues to serve as probes to characterize protein dynamics from picosecond to microsecond time-scales. We found that the native cellular environment only marginally influenced the inherent dynamics of MBP, but altered the molecular motion of the protein on a global level.

To facilitate the direct comparison of the native cellular conditions within OMVs to the buffer conditions of purified proteins, we isolated ILV-labeled MBP directly from the bacterial periplasm of the same cultures we harvested the MBP-enriched OMVs from (Fig 1A). Both samples yielded highly similar 2D NMR spectra of exceptional quality with only few resonances of differing chemical shifts (Fig. 1B). However, comparison to existing literature data showed, despite a global similarity, pronounced differences in the chemical shifts, which could be attributed to the presence of divalent (Ca^2+^ / Mg^2+^) ions in the buffer used in our experiments (Fig. S2A). These ions could not be omitted from our experimental set-up, since the stability of bacterial outer membranes and consequentially the stability of the OMVs used in our experiments demands the presence of low concentrations of divalent ions. In fact, absence of divalent ions results in substantial leakage of luminal contents from OMVs (not shown), whereas OMVs otherwise stably maintain their luminal content. To confirm the latter we re-collected OMVs by ultracentrifugation after running NMR experiments for two consecutive weeks and recorded 2D NMR spectra of the supernatant, which were virtually free of signals (Fig. S2B).

**Figure 1.**
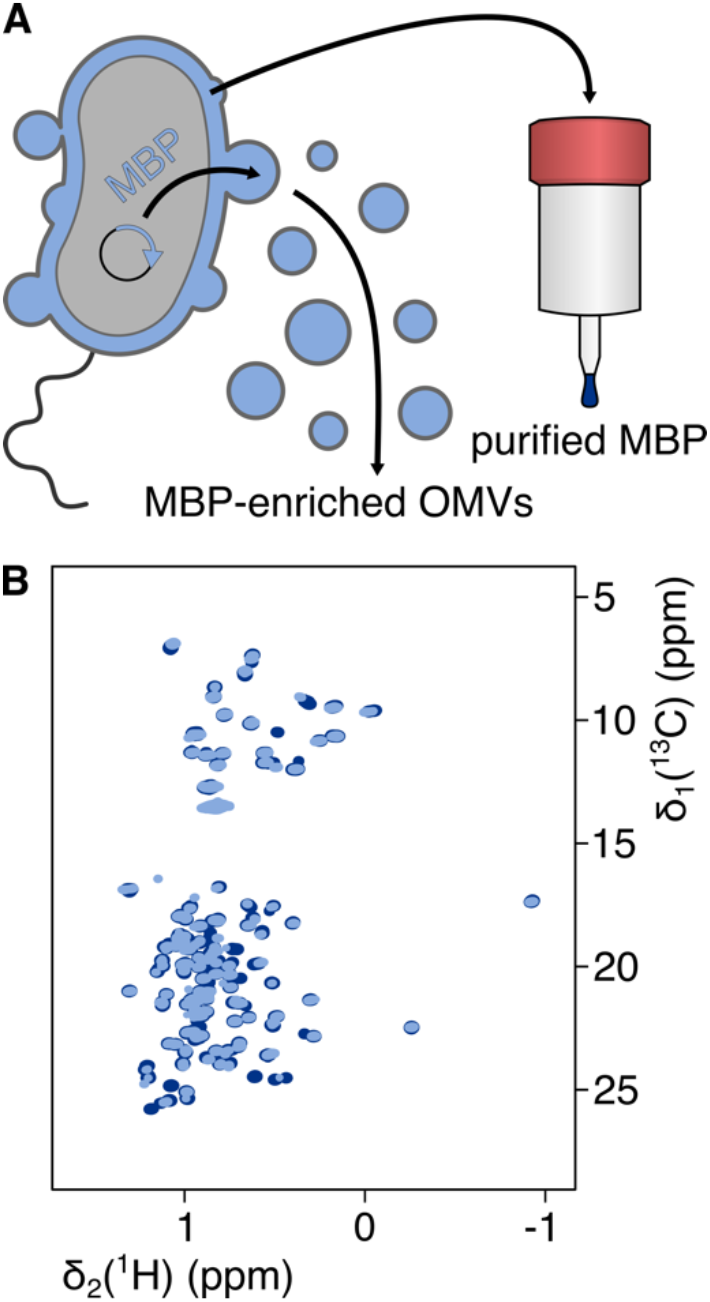
Experimental setup. (A) Sample preparation: Overexpression of the *MalE* gene in *Escherichia coli* results in enrichment of MBP in the bacterial periplasm. OMVs released from the bacteria are isolated and used for *in-situ* NMR spectroscopy. For comparison, MBP is purified from the periplasm of the same bacteria. (B) 2D [^13^C,^1^H]-NMR spectra of ILV-labeled MBP in OMVs (light blue) and purified MBP (dark blue).

Due to the differences caused by the buffer conditions, the majority of resonance assignments could not be readily transferred from existing literature data. We therefore resorted to obtaining resonance assignments based on available structures of MBP using methyl NOE data. To this end we recorded 3D C_Methyi_C_Methyi_H_Methyl_ NOESY spectra of ILV-labeled MBP purified from the bacterial periplasm (7). Recently, several automated approaches have been proposed to facilitate the assignment process (8–10). However, when applied to our data, the assignments generated by these methods were only in partial agreement with each other, necessitating laborious manual validation and adjustment of automatically obtained assignments. To link the intra-residue C^*δ*^ and C^*γ*^ methyl pairs of leucine and valine residues we recorded additional 3D HMBC-HMQC experiments (11). Resonances originating from valine residues were unambiguously identified using a V-labeled MBP sample (Fig. S3). Guided by existing and the automatically obtained partial assignments, we could confidently assign all resonances of ILV labeled MBP by matching experimentally determined NOE connectivity to available 3D structures (Table S1). Importantly, because the NOE network was in agreement with the connectivity expected based on the existing 3D structure of MBP (12), and because the differences between the MBP-enriched OMV sample and purified MBP were only minimal we can conclude that the tertiary structures of MBP in the native environment and purified MBP are highly similar.

Following resonance assignment we determined the influence of the native environment on the fast time-scale dynamics of MBP in the picosecond to nanosecond regime. To this end we determined the product S^2^_axis_.τ_C_ of the side-chain methyl order parameter and the rotational correlation time for MBP-enriched OMVs and purified MBP (13). Comparison of the values showed a striking overall similarity between the two conditions (Fig. S4), i.e. S^2^_axis_.τ_C_ values indicated side-chains of the equivalent residues to be more flexible (5 ns) or more rigid (20 ns) in the purified sample as well as in the native condition. Nevertheless, on average S^2^_axis_.τ_C_ values were ~30% higher in the OMV sample compared to purified MBP. To differentiate whether this difference can be attributed to either generally higher side-chain order parameter values or an increased rotational correlation time of MBP within the crowded lumen of OMVs we used the TRACT experiment to estimate τ_C_ of MBP in the OMV sample (Fig. S5). The estimated rotational correlation time of 20.7 ± 4.5 ns (in H_2_O based buffer) was substantially higher than values previously determined for purified MBP of 16.2 ± 0.7 ns (in H_2_O based buffer) (14). Using these values of τ_C_ to calculate the actual side-chain order parameters yielded highly similar distributions of S^2^_axis_ values, on average within a margin of ~1% and well within the experimental error (Fig. 2A). Nevertheless, despite the overall similarity for some residues we observed larger differences between the two conditions. Mapping these differences to the structure of MBP revealed that the residues that were affected most were generally located close to the solvent accessible surface of the protein (Fig. 2B), indicating a slight increase in S^2^_axis_ values of these residues. In contrast, residues within the folded core of MBP showed a mild trend towards lower S^2^_axis_ values. Taken together our data suggest that the inherent dynamics of the side-chain methyl groups of MBP are only mildly affected by the surrounding environment, whereas the global molecular motion appears to be strongly reduced within OMVs, as evidenced by the substantially increased rotational correlation time determined in our experiments.

**Figure 2.**
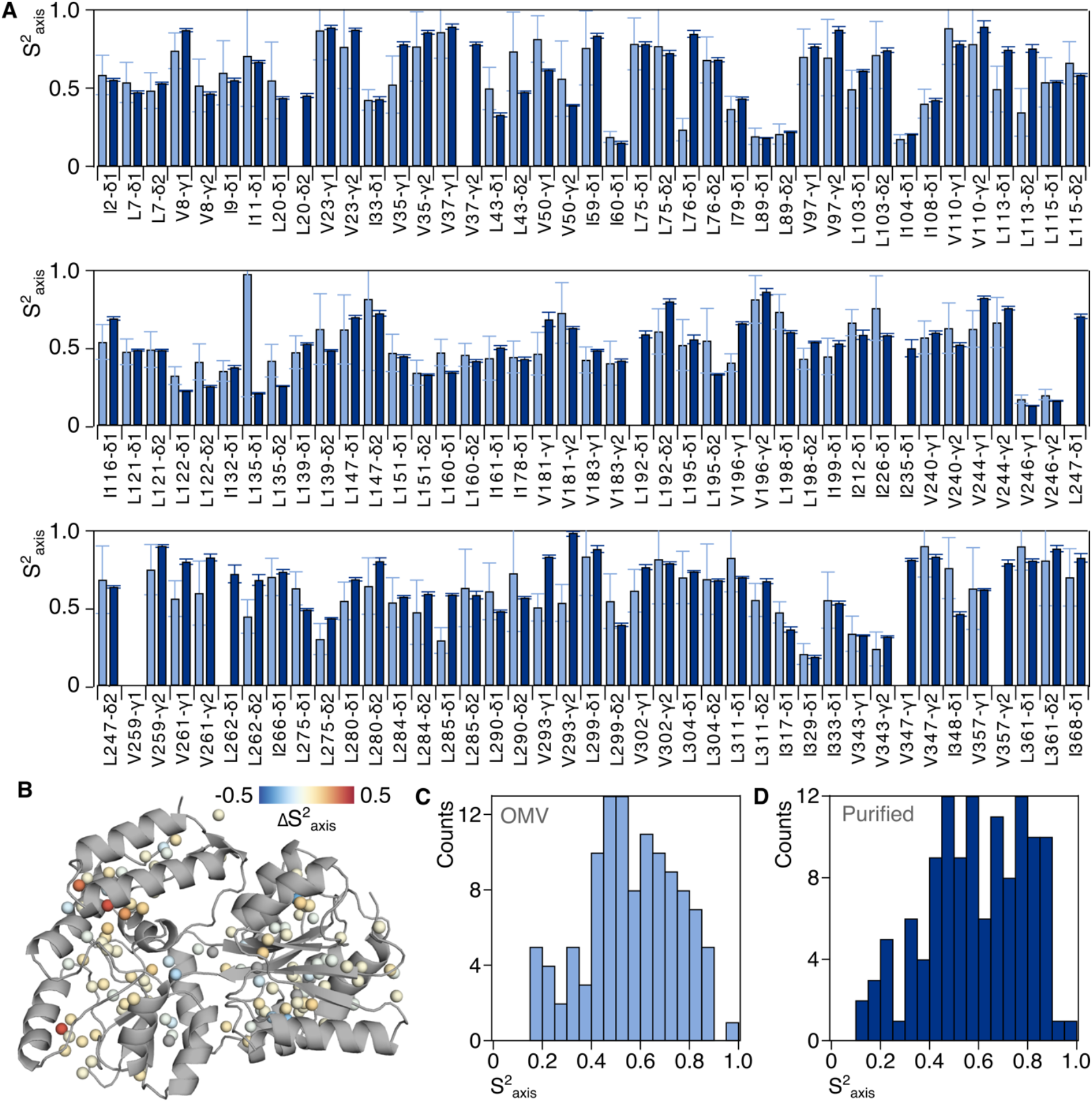
Fast dynamics of MBP side-chain methyl groups. (A) S^2^_axis_ values determined for MBP in OMVs (light blue) and purified MBP (dark blue) plotted against the sequence. (B) Difference of S^2^_axis_ values between the two conditions. C^δ^ and C^γ^ methyl groups are shown as spheres, difference is indicated by the color. (C, D) Histogram over determined S^2^_axis_ values for MBP in OMVs (C) and purified MBP (D).

To probe whether the native environment imposed differences on the slow dynamics of MBP in the microsecond to millisecond regime we subsequently utilized multiple quantum Carr-Purcell-Meiboom-Gill (CPMG) relaxation dispersion experiments (15). While we obtained generally flat profiles for the majority of methyl groups, some appeared to be involved in conformational exchange processes, indicated by non-flat dispersion profiles. Importantly, as previously observed with the order parameters, the overall distribution of ΔR_2,eff_ values was highly similar between the two conditions (Fig. 3A), suggesting that the inherent dynamics of MBP also on the slower timescale are largely unaffected by the surrounding environment. As previously observed, the largest differences in ΔR_2,eff_ between the two conditions were observed for methyl groups near the solvent accessible surface of MBP (Fig. 3B).

**Figure 3.**
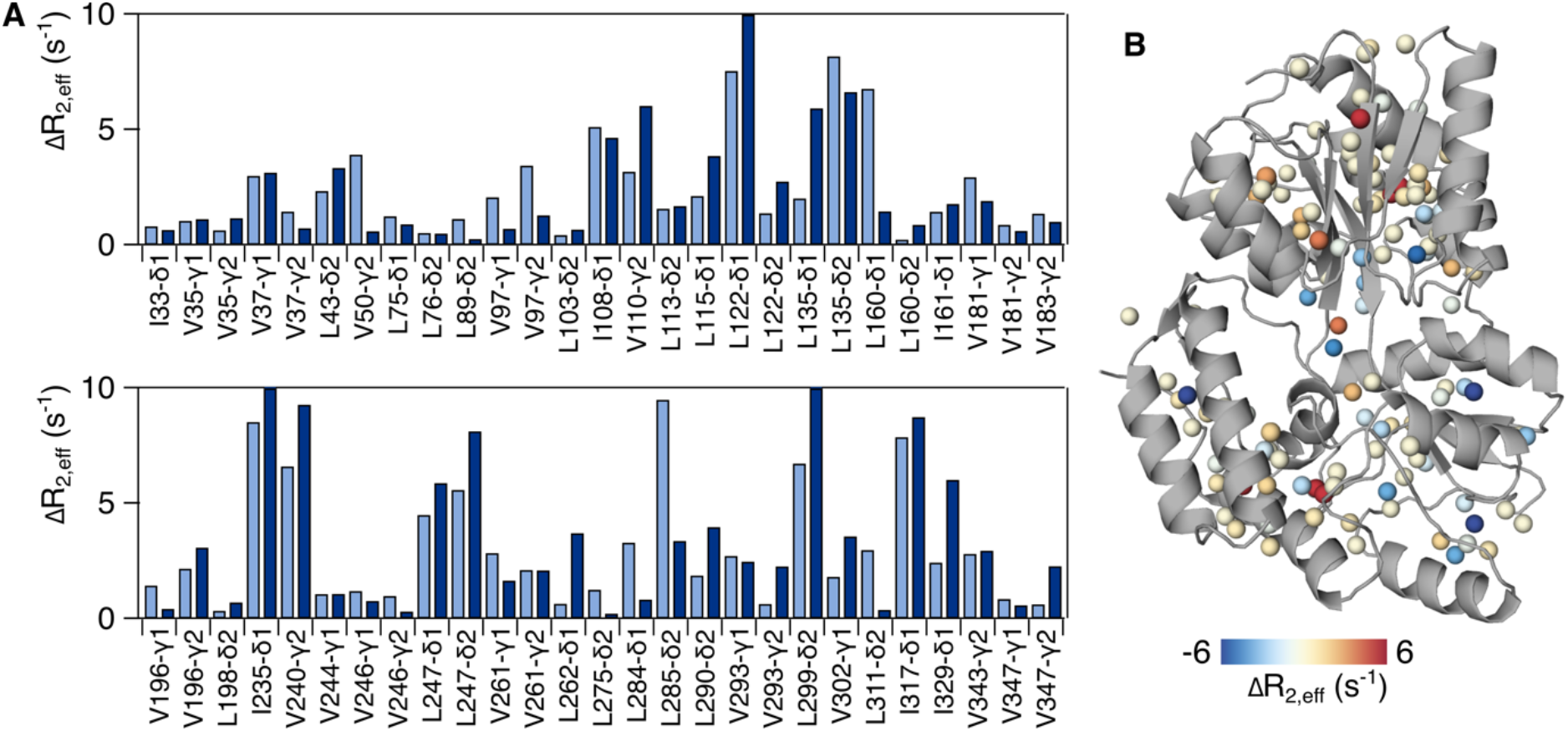
Slow dynamics of MBP side-chain methyl groups. (A) ΔR_2,eff_ values determined for MBP in OMVs (light blue) and purified MBP (dark blue) plotted against the sequence. Only residues with ΔR_2,eff_ > 0.5 s^-1^ are shown. (B) Difference of ΔR_2,eff_ values between the two conditions. C^δ^ and C^γ^ methyl groups are shown as spheres, difference is indicated by the color.

In conclusion, our experiments clearly demonstrate the feasibility of using OMVs to determine the dynamics of periplasmic proteins at exceptional detail within their native cellular environment by exploiting the high sensitivity of methyl-TROSY based NMR spectroscopy methods. Thereby our data suggest that across timescales the dynamics of MBP in the native lumen of OMVs closely resemble the dynamics of purified MBP, despite overall slower molecular tumbling. The latter can be attributed to the high periplasmic protein concentration and the resulting increased viscosity within the crowded native environment. We can therefore conclude that the inherent dynamics of the “Teflon-protein” MBP are largely immune to the molecular environment.

## Acknowledgements

The Swedish NMR Centre of the University of Gothenburg is acknowledged for spectrometer time. J.T. was supported by an EMBO Long-Term Fellowship (ALTF 413-2018). B.M.B. gratefully acknowledges an EMBO Young Investigator Fellowship as well as funding from the Swedish Research Council (Starting Grant 2016-04721; Consolidator Grant 2020-00466), the Swedish Cancer Foundation (2019-0415), and the Knut och Alice Wallenberg Foundation through a Wallenberg Academy Fellowship (2016.0163) as well as through the Wallenberg Centre for Molecular and Translational Medicine, University of Gothenburg, Sweden.

## Supporting Material

**Supplementary Figure 1.**
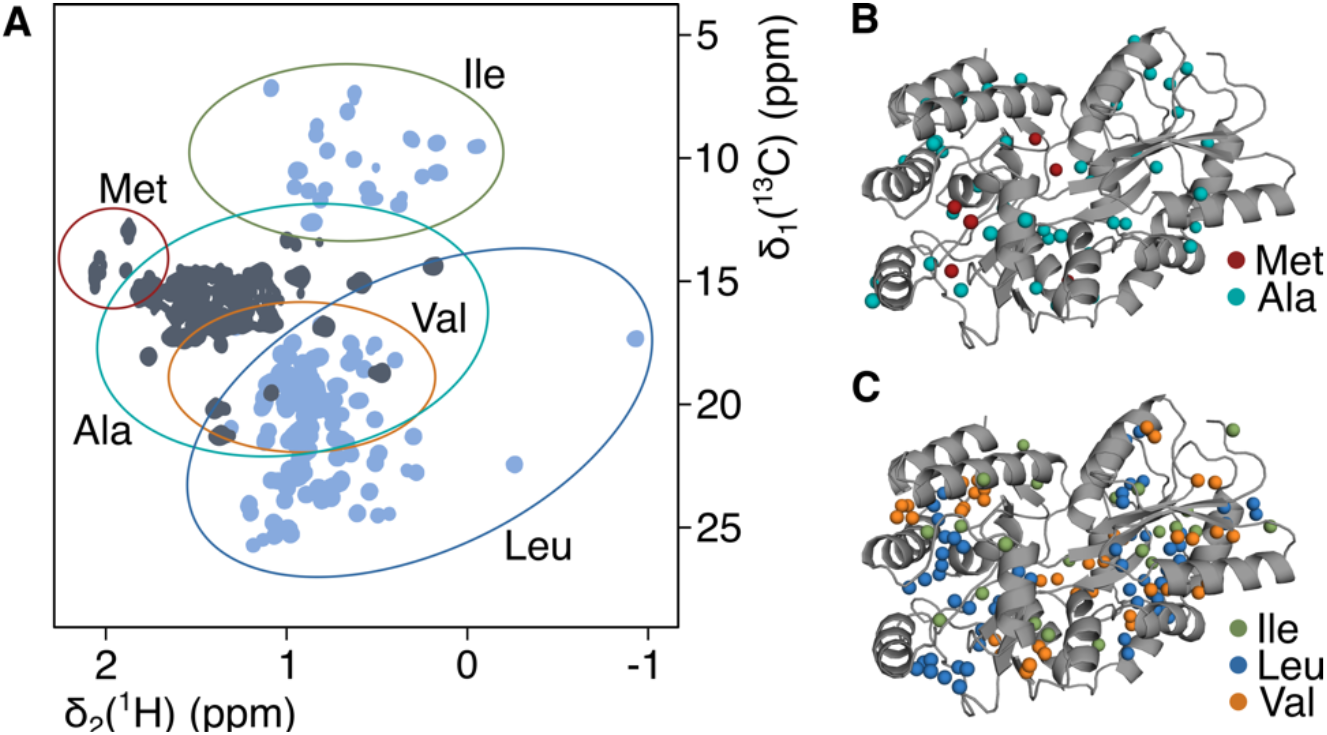
Side-chain methyl labeling of MBP. (A) 2D [^13^C,^1^H]-NMR spectra of ILV-labeled MBP (light blue) and MA-labeled MBP (grey). Due to pronounced overlap of alanine C^β^ resonances in one densely clustered region we relied on the ILV labeling scheme for further experiments. (B, C) Localization of side-chain methyl groups in the 3D structure of MBP. Methyl groups are shown as spheres as indicated by the color legends.

**Supplementary Figure 2.**
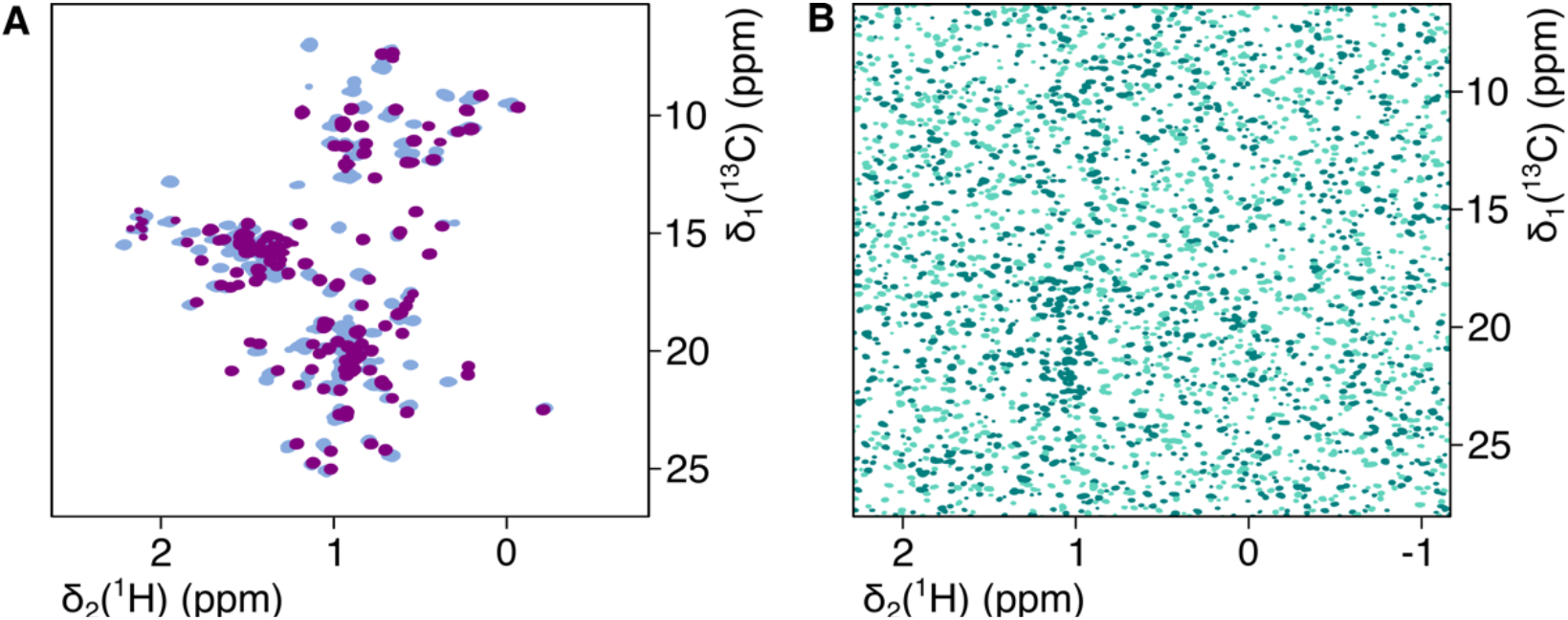
Effect of Buffer Conditions. (A) 2D [^13^C,^1^H]-NMR spectra of ILV-(*proS*)-labeled MBP in NMR buffer (see Methods) containing Ca^2+^ and Mg^2+^ (light blue) and in absence of divalent ions (purple). (B) 2D [^13^C,^1^H]-NMR spectrum of supernatant following the removal of OMVs by centrifugation.

**Supplementary Figure 3.**
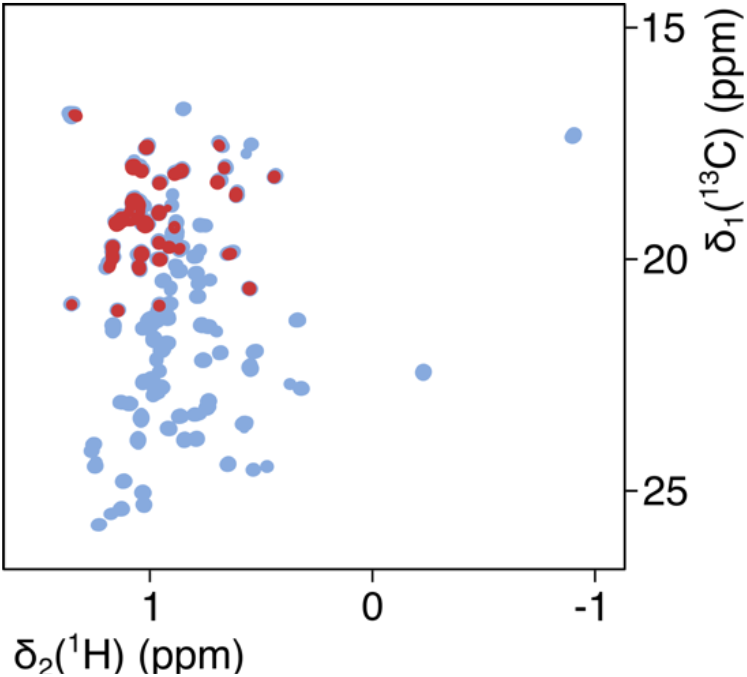
Identification of valine resonances. 2D [^13^C,^1^H]-NMR spectra of ILV-labeled MBP (light blue) and valine-labeled MBP (red).

**Table 1.**
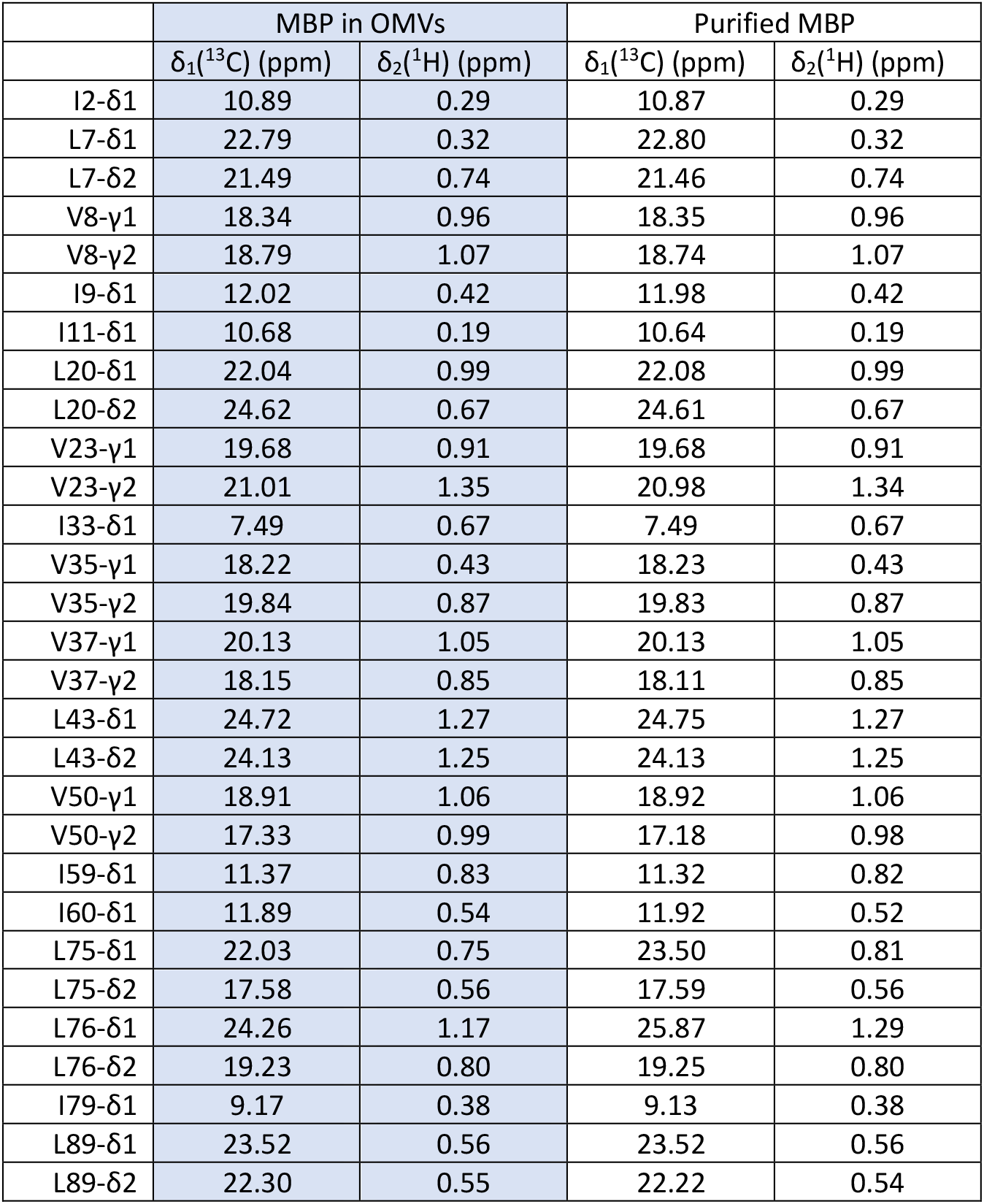
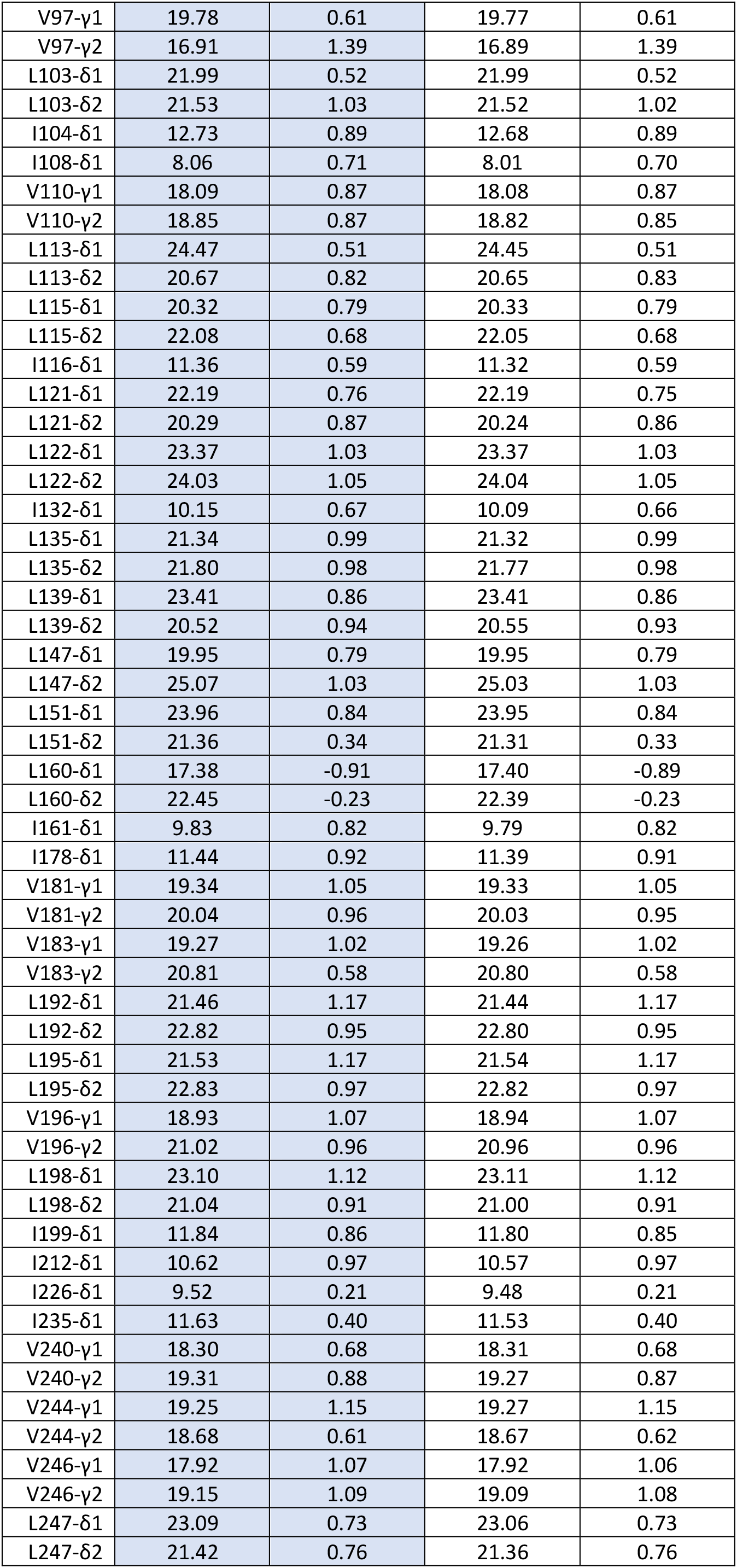
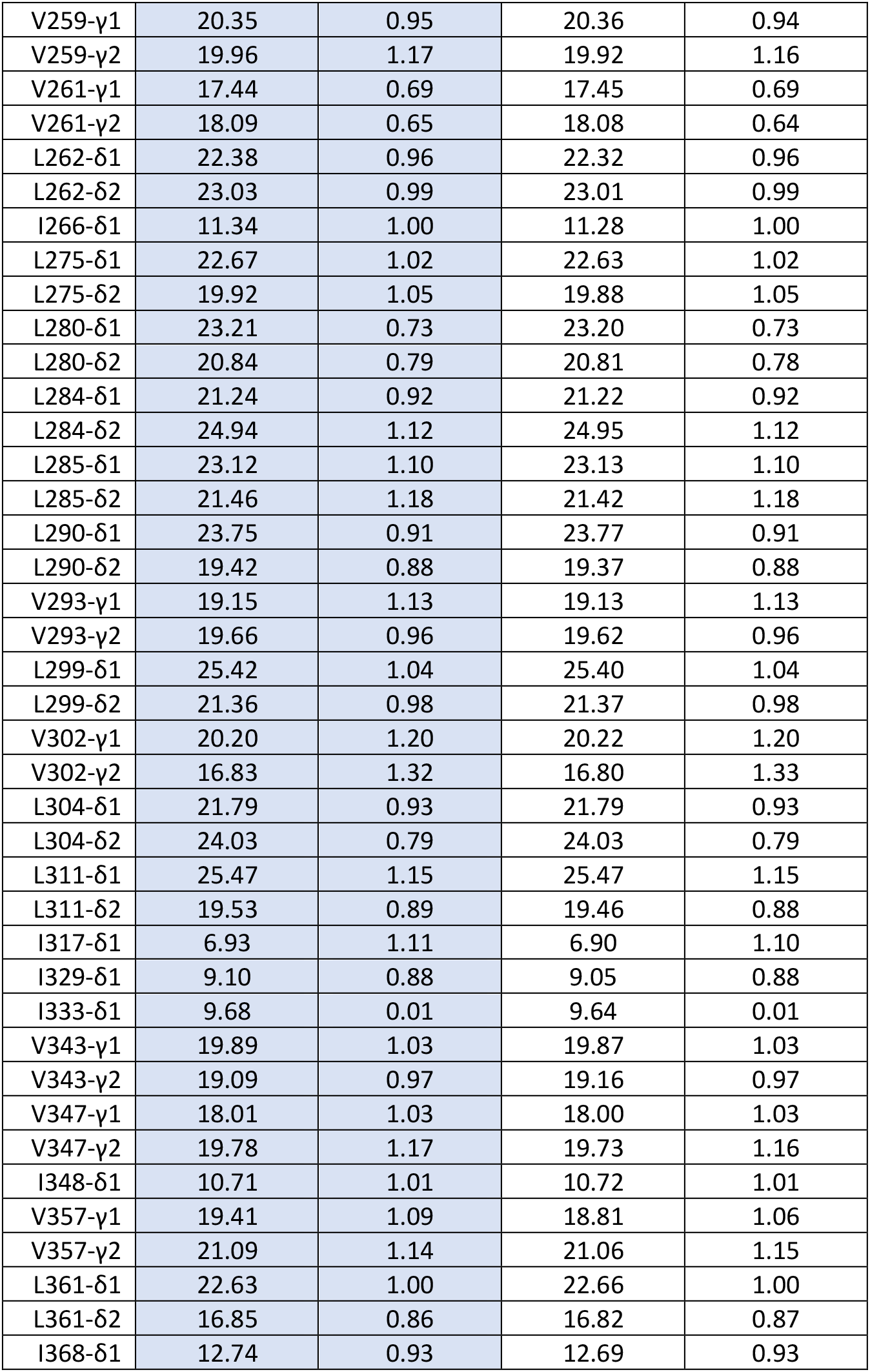
Resonance assignment of MBP in OMVs and purified MBP.

**Supplementary Figure 4.**
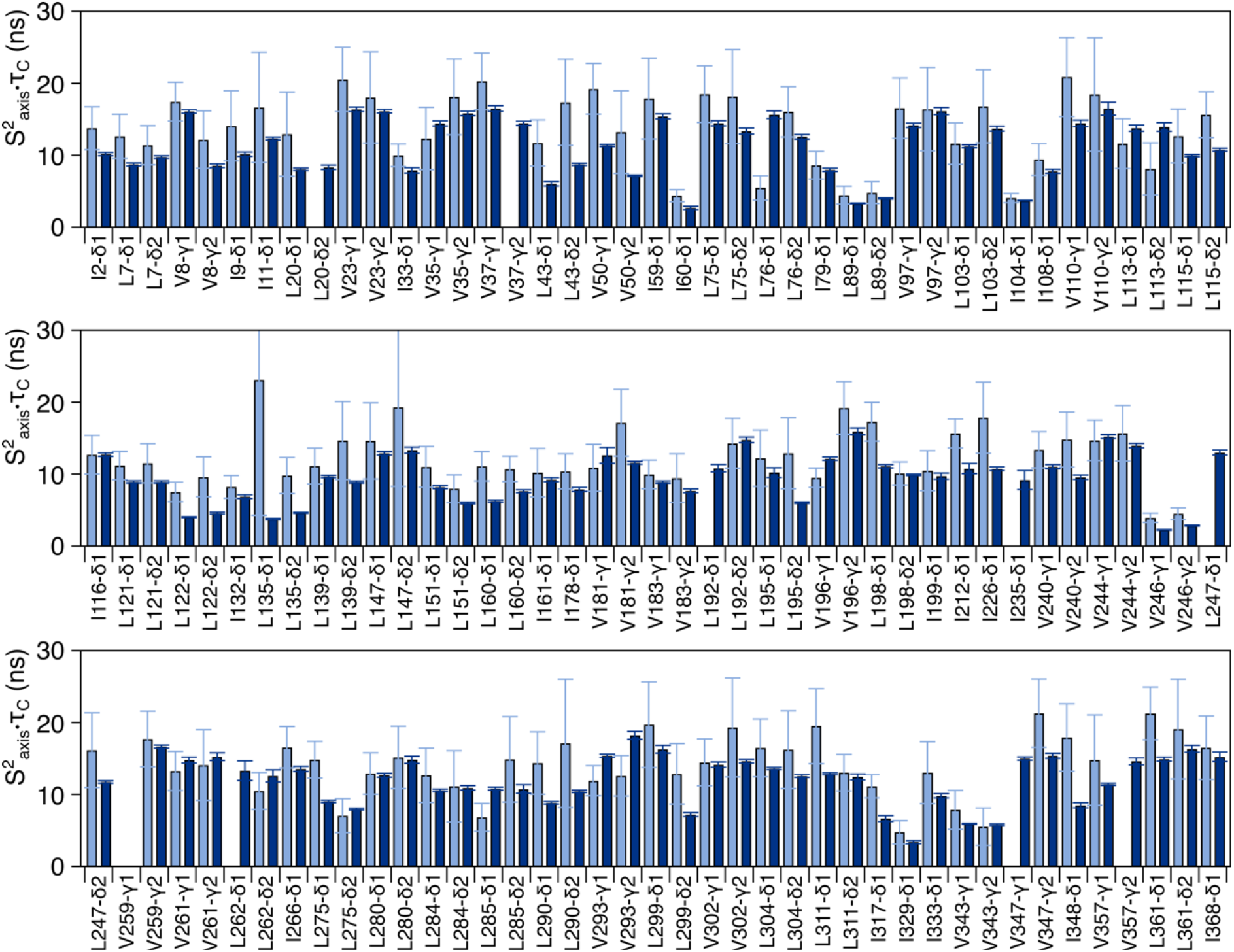
Fast dynamics of MBP side-chain methyl groups. S^2^_axis_.τ_C_ values determined for MBP in OMVs (light blue) and purified MBP (dark blue) plotted against the sequence.

**Supplementary Figure 5.**
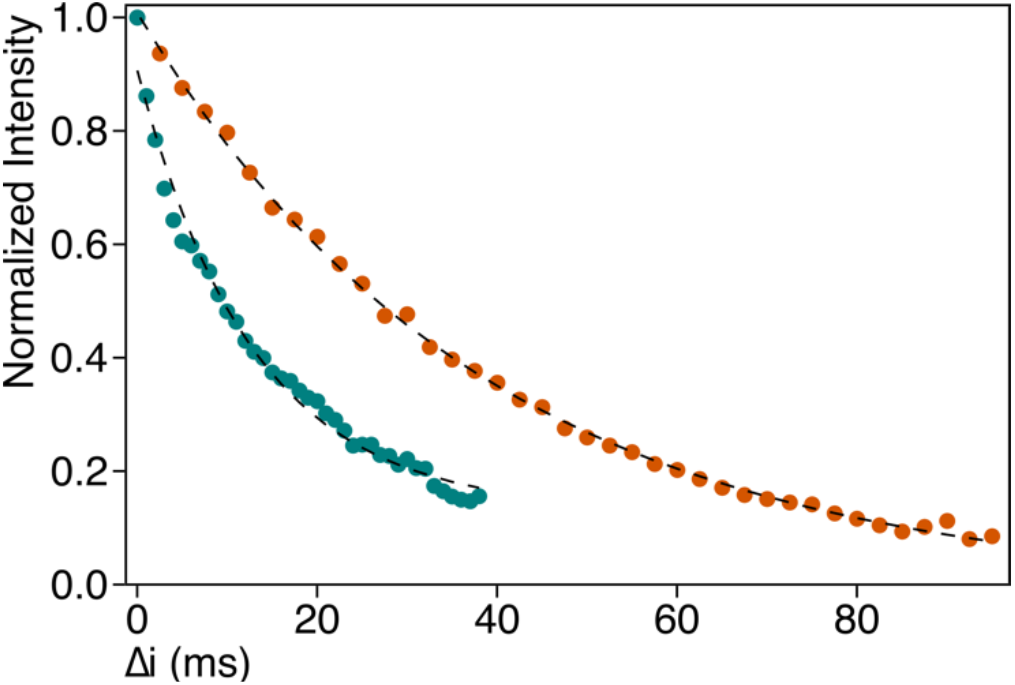
Determination of the rotational correlation time τ_C_ of MBP in OMVs. Normalized integral values are plotted against the relaxation period Δi for the α-(orange) and β-spin state (teal). Dashed lines show exponential fit functions.

## Methods

### Preparation of MBP-enriched OMVs

MBP-enriched OMVs form *E. coli* strain BL21(DE3)ΔompA (1) were prepared using plasmid pY271 (2) as described. Bacterial cultures were inoculated at a ratio of 1/100 from liquid cultures and grown overnight in 50 mL M9 minimal medium (3) containing [^15^N]H_4_Cl and [^12^C_6_D_12_]O_6_ D-Glucose and 99.8% D_2_O, supplemented with 100 μg mL^-1^ Ampicillin in baffled 1 L Erlenmeyer flasks at 37 °C. Overnight cultures were transferred to 450 mL fresh medium and grown in baffled 5 L Erlenmeyer flasks at 37 °C. When cultures reached an optical density of OD_600_ ≈ 0.35 precursors ketoisovalerate and ketobutyrate (Merck) were added to a final concentration of 85 mg L^-1^ and 50 mg L^-1^, respectively, to achieve selective labeling of either one (*proS*: DLAM-LV^*proS*^-kit from NMR Bio) or both of the C^δ1/2^ and C^γ1/2^ methyl groups of isoleucine, leucine and valine residues. For selective labelling of the C^b^ and C^ε^ methyl groups of alanine and methionine (Cambridge Isotope Labs), respectively, alanine and methionine were added to a final concentration of 50 mg L^-1^. For the valine-labeled sample ketoisovalerate was added to a final concentration of 40 mg L^-1^ together with deuterated leucine (Merck) to a final concentration of 25 mg L^-1^. When cultures reached an optical density of OD_600_ ≈ 0.4 expression of the *malE* gene was induced by addition of 0.25 mM IPTG. After 5 h bacteria were removed by centrifugation and processed further as described below. Culture supernatants were filtered through 0.45 μm PVDF filter units (Merck Millipore). OMVs were collected by centrifugation (38’400 xg for 2 h), resuspended in 25 mL NMR buffer (Dulbecco’s PBS with added salts: 8.1 mM Na_2_HPO_4_, 1.5 mM KH_2_PO_4_, 2.7 mM KCl, 136.9 mM NaCl, 0.9 mM CaCl_2_, 0.5 mM MgCl_2_, 1 mM Maltose, pH 7.4; prepared either in 99.8% D_2_O or in 90% H_2_O / 10% D_2_O (4)), filtered using 0.45 μm PES syringe filters and recollected by ultracentrifugation (74’500 xg for 45 min). OMVs were then resuspended in a final volume of 180 μL NMR buffer.

### Purification of periplasmic MBP

Bacterial pellets were resuspended in 5 mL buffer (200 mM Tris, 1 M Sucrose, 1 mM EDTA, pH 8) and 5 mg Lysozyme were added. Following incubation at room temperature for 5 min 20 ml H_2_O were added. The cell suspension was mixed by inversion and incubated on ice for 30 min to allow spheroplast formation. Cells were then pelleted at 200′000 xg for 30 min. The cleared supernatant was dialyzed twice against 2 L binding buffer (20 mM Tris, 200 mM NaCl, 1 mM EDTA, pH 7.4) and applied to a 5 mL MBPTrap HP column (GE Healthcare) preequilibrated with binding buffer. The column was then washed with 50 mL binding buffer. MBP was eluted 25 mL elution buffer (10 mM Maltose, 20 mM Tris, 200 mM NaCl, 1 mM EDTA, pH 7.4) in 1 mL fractions. Fractions containing MBP were pooled, concentrated using centrifugal concentrators (Vivaspin 6, 10 kDa MWCO), and the buffer was exchanged to NMR buffer by repeated dilution and concentration.

### NMR Spectroscopy

NMR measurements were performed on Bruker AscendII 700 MHz, 800 MHz, and 900 MHz spectrometers running Topspin 3.6 and equipped with cryogenically cooled triple-resonance probes. All NMR-experiments were performed in D_2_O or H_2_O based NMR buffer as stated and at a temperature of 310 K. The following pulse schemes were used: 2D SOFAST HMQC (5), 3D C_Methyl_C_Methyl_H_Methyl_ SOFAST NOESY-HMQC with mixing times of 50 ms and 300 ms (6), 3D HMBC-HMQC (7). For the determination of side-chain methyl order parameters single- and triple-quantum ^1^H–^13^C relaxation experiments were used with delay times of 1.1, 2, 4, 8, 12,…, 40 ms (8). TRACT experiments were recorded using relaxation periods Δi of 2.5, 5, 7.5,…, 95 ms for the a- and 1, 2, 3,…, 38 ms for the β-spin state (9). Multiple quantum relaxation dispersion experiments were recorded using constant time relaxation periods of 40 ms and CPMG frequencies from 67 to 1,000 Hz (10).

### Data processing and analysis

NMR data were processed with TopSpin 4.0 (Bruker BioSpin), NMRpipe (11), and mddnmr (12). Resonance assignment was done in an assisted manual approach using CCPN (13) in combination with the MAUS (14), Methyl-FLYA (15), and MAGIC (16) algorithms. Peak intensities for the determination of methyl order parameters and multiple quantum relaxation dispersion experiments were determined using PINT(17). For order parameter determination ratios of the peak intensities of the single- and triple-quantum experiments were fitted using the equation:

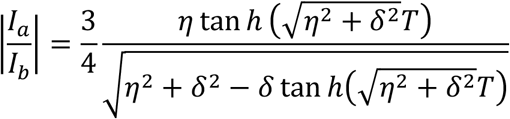

and S^2^_axis_ values were determined using equation

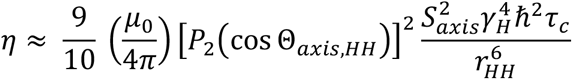

as described elsewhere (8). The rotational correlation time τ_C_ of MBP in OMVs was determined using exponential fit and equation τ_C_ = ((τ_2_-τ_1_)/(5·ω_N_)·1e^9^ as described (9). Multiple quantum methyl relaxation dispersion data were fitted to a two-site exchange model using ChemEx (available at https://github.com/gbouvignies/chemex/releases).

